# Dietary Proanthocyanidins Exert Localized Immunomodulatory Effects in Porcine Pulmonary and Gastrointestinal Tissues during *Ascaris suum*-induced Type 2 inflammation

**DOI:** 10.1101/2021.10.12.464117

**Authors:** Audrey Inge Schytz Andersen-Civil, Laura J. Myhill, Nilay Büdeyri Gökgöz, Marica T. Engström, Helena Mejer, Wayne E. Zeller, Juha-Pekka Salminen, Lukasz Krych, Charlotte Lauridsen, Dennis S. Nielsen, Stig M. Thamsborg, Andrew R. Williams

## Abstract

Bioactive dietary components may considerably influence intestinal health and resistance to enteric disease. Proanthocyanidins (PAC) are dietary polyphenols with putative health-promoting activity that have been increasingly studied for their anti-inflammatory and immunomodulatory effects. However, whether dietary PAC can regulate type-2 immune function and inflammation at mucosal surfaces remains unclear. Here, we investigated whether diets supplemented with purified PAC modulated pulmonary and intestinal mucosal immune responses during infection with the helminth parasite *Ascaris suum* in pigs. *A. suum* infection induced a type 2-biased immune response in lung and intestinal tissues, characterized by pulmonary granulocytosis, increased Th2/Th1 T cell ratios in tracheal-bronchial lymph nodes, intestinal eosinophilia, and modulation of genes involved in mucosal barrier function and immunity. We observed that PAC had only minor effects on pulmonary immune responses, regardless of concurrent *A. suum* infection. However, RNA-sequencing of intestinal tissues revealed that dietary PAC significantly enhanced transcriptional responses related to immune function, antioxidant responses, and cellular stress activity, both in uninfected and *A. suum*-infected animals. *A. suum* infection and dietary PAC both induced distinct changes in gut microbiota composition, primarily in the jejunum and colon, respectively. Notably, PAC substantially increased *Limosilactobacillus reuteri* abundance in the colon of both naïve and *A. suum*-infected animals. Thus, dietary PAC may have distinct beneficial effects on intestinal health during infection with mucosal pathogens, whilst having limited activity to modulate naturally-induced type-2 pulmonary inflammation. Our results shed further light on the mechanisms underlying the health-promoting properties of PAC-rich foods, and may aid in the design of novel dietary supplements to regulate mucosal inflammatory responses in the gastrointestinal tract.

## Introduction

Effective immune function is essential for maintenance of health and tissue homeostasis. The role of diet in regulating immunity and inflammation at mucosal barrier surfaces has been well-established, and immunomodulatory dietary components have therefore gained tremendous attention in scientific research in recent years. Polyphenols, terpenoids, and carotenoids are examples of three central groups of phytonutrients, which have been extensively studied for their beneficial impact on health and disease^1–4^. Proanthocyanidins (PAC) are a type of polyphenol, commonly found in a plant-based diet, which have characteristic chemical structures with known anti-oxidant and anti-inflammatory properties^5^.

Numerous studies have demonstrated that PAC play an important role in the regulation of immune function and may offer therapeutic potential towards inflammatory intestinal diseases. PAC exert strong antioxidant and anti-inflammatory effects in cellular models^6, 7^, and are able to modulate various physiological parameters when consumed as part of the diet, such as increasing mucus production in human patients with ulcerative colitis, and alleviating inflammation in TNBS-induced colitis by inhibiting NF-kB signaling pathways^8, 9^. Another consistent outcome of dietary PAC supplementation is changes in gut microbiota (GM) composition, which has been observed in both murine and porcine models, with some evidence also from human studies^10–14^. PAC have been shown to increase the abundance of Lactobacilli and *Bifidobacterium* species, which are commonly associated with a healthy gut environment, as well as increasing levels of faecal short chain fatty acids (SCFA) such as propionic acid^11, 15, 16^. Thus, the immunomodulatory effects of PAC may be caused by direct interactions with immune cells and/or indirect modulation of immune responses as a result of PAC-induced alteration of the GM^17^. These bioactive effects on intestinal immune cells or the GM may have significant implications for inflammation in the gut, as well as at distant sites, such as the lungs, with increasing evidence suggesting a gut-lung axis and clear connection between gut function, the microbiome, and lung homeostasis^18^.

Parasitic worm (helminth) infections are widespread in humans and animals worldwide and cause substantial morbidity^19, 20^. The characteristics of immune responses during helminth infection include a strongly Th2 polarized immune response, characterized by eosinophilia, mastocytosis and increased production of IL-4, IL-13 and other type-2 cytokines^21^. Thus, helminth infection models offer a valuable opportunity to assess how different dietary interventions can promote resistance to parasitic infection, as well as modulating type-2 responses which play a significant role in pathologies such as allergies. In mice, PAC may regulate allergen-induced type-2 inflammation in the lungs by decreasing the expression of IL-4, IL-5 and IL-13^22^. However, the ability of PAC to modulate pathogen-induced type-2 mucosal immune responses, such as those induced by tissue-invasive helminths, has not been examined in detail. Such studies may shed light on the interactions between PAC-rich diets and immunity to helminth infection, and other type-2 driven pathologies, such as asthma and ulcerative colitis.

Pigs are a highly translatable large-animal model for humans, due to the anatomical and immunological similarities between humans and swine. The porcine roundworm *Ascaris suum* is widely prevalent in pig farms globally and closely related to *A. lumbricoides*, which is the most common helminth in humans^23^. After infection, *A. suum* larvae have a complex migratory path, which includes migration through the liver and lungs before returning to the small intestine^24^. At each of these anatomical sites, the migratory larvae cause strong inflammatory reactions. Studies in both the natural porcine host and murine models have shown that larvae elicit significant levels of type-2 (e.g. IL-5), but also type-1/17 (IL-6 and TNFα) cytokines as they migrate in the liver, lungs and gut^25, 26^. Furthermore, *A. suum* infection has been shown to increase the susceptibility to bacterial lung infections in both mice and pigs^27, 28^.

The life cycle of *A. suum*, with larvae relocating between the intestines, the liver and the lungs, offers numerous sites of interaction with the immune system. In this respect, *A. suum* infections in pigs have proved useful for assessing the effects of different dietary components on type-2 immune function in both the gut and the respiratory tract. For example, studies in this model have shown that dietary retinoic acid can enhance *A. suum*-induced pulmonary eosinophilia, whilst treatment with probiotics, such as *Lactobacillus rhamnosus* GG, can suppress the prototypical type-2 response in lymph nodes draining the lungs during infection^29, 30^. However, studies on the effects of concomitant PAC-supplementation and helminth infection are scant. Here, we explored the effect of dietary PAC on host immune function in *A. suum*-infected pigs. We examined the impact of PAC on systemic immune parameters, inflammatory and immune reactions at the mucosal barrier of both the lung and the intestinal tract, and infection- and dietary-induced changes in the gut microbiota. Thus, the aim was to investigate how PAC consumption may modulate a naturally-induced type-2 mucosal response in multiple tissue sites.

## Material and Methods

### Proanthocyanidins and diets

The PAC used for this study were from a standardized grape seed extract (Bulk Powders, Denmark). Based on LC-DAD-MS and LC-DAD-MS/MS analyses^31, 32^, the PAC purity of the extract was >95%. Further analysis of the PAC showed that they were composed of 99% procyanidin oligomers and polymers, with a mean degree of polymerization of 4.2. The basal diet (NAG, Denmark) was based on ground wheat and barley and was formulated to provide 16.2% crude protein (**Supplementary Table 1**). Pigs received either the basal diet or the basal diet supplemented with 1% PAC. Feed intake was adjusted for body weight throughout the experiment and was calculated to provide the PAC-supplemented pigs with approximately 300 mg PAC/kg BW. All pigs were weighed weekly and they were monitored and fed twice daily at 8:00 in the morning and 15:00 in the afternoon, with access to water *ad libitum*.

### Pig experiment

9-weeks old pigs were selected from a Specific Pathogen Free (SPF) Danish farm with no history of helminth infection. On arrival, pigs were confirmed free of helminth-infection by fecal egg count and negative by serology for *A. suum*. Pigs were vaccinated (p.o.) against *Lawsonia intracellularis* 4.5 weeks prior to the start of the experiment (ENTERISOL^®^ ILEITIS VET., Boehringer Ingelheim). A total 24 pigs (Duroc/Danish Landrace/Yorkshire; 12 castrated males and 12 females) were randomly distributed into four treatment groups that were balanced for sex and initial bodyweight. Bodyweights were recorded weekly (**Supplementary Figure 1**). Each of the four groups were housed in two pens consisting of three pigs each. From day 1 of the experiment, 12 pigs were fed the basal diet and 12 the PAC-supplemented diet. At day 14, half the pigs in each group were inoculated with 5000 embryonated *A. suum* eggs by gastric intubation (**Supplementary Figure 2**). Pigs were euthanized at day 28 of the experiment, i.e. day 14 post-infection, (p.i.) by captive bolt pistol stunning followed by exsanguination. Throughout the study, weekly blood and fecal samples were taken. Blood was collected by venipuncture of the jugular vein and serum separated and frozen at −80 °C. At necropsy, the entire small intestine was removed and processed for *A. suum* larval counts using a modified agar-gel technique^33^. Worm burdens were assessed by manual enumeration using a dissection microscope of blinded samples conserved in 70% ethanol. Digesta samples were collected from the proximal colon and cooled on ice before transfer to – 80°C storage. Small pieces (1cm^3^) of lung (right cranial lobe) and mid-jejunal tissue were preserved in RNAlater. A further piece of jejunal tissue was also collected for histology using BiopSafe^®^ Biopsy Sample System (Merit Medical). Histology slides were stained with hematoxylin & eosin, and eosinophils were enumerated by blinded microscopy.

### Broncho-Alveolar Lavage

Broncho-alveolar lavage (BAL) was performed at necropsy by introducing 500 ml PBS into the lungs to recover BAL cells from both lung lobes. The BAL fluid was filtered through 2-layer fine gauze sheets into clean 50 mL centrifuge tubes, and stored at room temperature (RT) until further processing. The recovered cell suspensions underwent a series of washing with HBSS and centrifugation. To remove red blood cell (RBC) contamination the cell suspension was incubated for 5 minutes at RT in RBC lysis buffer (Sigma-Aldrich). Finally, the cells were resuspended in 5 mL RPMI-1640 media supplemented with 10% foetal bovine serum and penicillin/streptomycin. Cells were enumerated and either used for flow cytometry (see below) or plated out on 48-well plates at a concentration of 1.2 x 10^5^ cells/well and incubated overnight (37 °C, 5% CO2). The next day, cells were stimulated with either excretory/secretory (E/S) products from either *A. suum* or *Trichuris suis*, or lipopolysaccharide (LPS; 500 ng/mL) for 24 hours. Following stimulation, the supernatant was collected and stored at – 20 °C before further analysis by ELISA.

### DNA extraction and 16S rRNA gene amplicon sequencing

DNA from small and large intestine samples was extracted using Bead-Beat Micro AX Gravity Kit (A&A Biotechnology, Gdynia, Poland) as per manufacturer’s instructions. The DNA purity and concentration were determined by NanoDrop 1000 Spectrophotometer (Thermo Fisher Scientific, USA) and Qubit™ 1x dsDNA high sensitivity kit on Varioskan Flash (Thermo Fisher Scientific, USA), respectively.

A 16S rRNA gene amplicon library was constructed by amplifying the 16S rRNA gene with unique molecular identifier (UMI) containing multiple forward and reverse primers (**Supplementary Table 2**). PCR conditions for the amplification were as follows: 95°C for 5 min, 2 cycles of 95°C for 20 s, 48°C for 30 s, 65°C for 10 s, 72°C for 45 s, and a final extension at 72°C for 4 min. A second PCR step was then performed to barcode PCR amplicons with the following conditions: 95°C for 2 min followed by 33 cycles of 95°C for 20 s, 55°C for 20 s, 72°C for 40 s, and a final extension at 72°C for 4 min. After each PCR reaction, PCR amplicons were cleaned up using SpeedBeads™ magnetic carboxylate (obtained from Sigma Aldrich). The size of barcoded PCR products (approximately 1500 bp) was checked by 1.5% agarose gel electrophoresis.

A sequencing library from pooled barcoded PCR products were prepared by following the ligation sequencing kit SQK-LSK110 (Oxford Nanopore Technologies, Oxford, UK) protocol. Next, prepared library was sequenced by Oxford Nanopore GridIONX5 sequencing platform as described in manufacturer’s protocol (https://nanoporetech.com/products/gridion). Sequencing was run until there was no longer active pores.

### Data analysis workflow for 16S rRNA gene sequencing

Nanopore sequencing software GridION version 21.02.5 (https://nanoporetech.com) was used for data collection. Base calling and demultiplexing of sequencing data were performed by ONT’s Guppy version 4.5.2 (https://nanoporetech.com). Nanofilt version 2.7.1^34^ was then used for filtering and trimming of demultiplexed sequences. Briefly, data were filtered on a minimum 1000 and maximum 1600 reads with a minimum average read quality score of 8. After filtering, 15 nucleotides were trimmed from start and end of reads. Taxonomy assignment was achieved by using parallel_assign_taxonomy_uclust.py script of Quantitative Insights into Microbial Ecology (Qiime) 1 version 1.8.0^35^. Greengenes database version 13.8 was used as a reference database^36^. The reads classifications did not include UMI correction due to low coverage of UMI clusters.

Qiime 2 version 2020.6.0^37^ was used to set rarefaction depth to 5000 reads per sample. Sample reads below 5000 were removed from the analysis; a total of 42 samples were included for microbiome analysis (n=19 for small intestine samples and n= 23 for large intestine samples). Normalized data were then processed in RStudio version 1.3.1073 using R version 4.0.2 and R packages phyloseq^38^, vegan^39^, tidyverse^40^, ggpubr^41^, reshape2^42^ and viridis^43^.

Raw 16S rRNA sequence data is available at Sequence Read Archive (www.ncbi.nlm.nih.gov/sra/) under accession number PRJNA753018.

### Measurement of gut microbial metabolites

Short-chain fatty acids and DL-lactic acid were analyzed in colonic digesta samples by GC-MS as previously described^44^.

### Flow-cytometry

Tracheal-bronchial lymph nodes were collected from the bifurcature and stored on ice in FBS supplemented RPMI until further processing. Single-cell suspensions were prepared by passing lymph nodes through a 70 μm cell-strainer. Cells were washed and stained with the following antibodies: mouse anti-pig CD3-FITC (clone BB23-8E6-8C8; BD Biosciences); mouse anti-pig CD4-PE-Cy7 (clone 74-12-4; BD Biosciences); mouse anti-human T-bet-APC (clone 4B10; BioLegend); mouse anti-human GATA3-PE (clone TWAJ; Invitrogen). BAL cells were collected as described above, and stained with mouse anti-pig granulocytes-Alexa Fluor647 (clone 2B2; Bio-Rad) and mouse anti-pig CD203a-FITC (clone PM18-7; Bio-Rad). Granulocytes were defined as 2B2^+^CD203a^-^. For all stainings isotype controls were included and gates were set using FMO controls. Data was acquired on an Accuri C6 flow cytometer (BD Biosciences) and analyzed using C6 software.

### Enzyme-linked immunosorbent assay

IL-1β and TNFα concentrations in alveolar macrophage supernatant and CRP levels in serum were analyzed using commercial ELISA kits (Duosets; R and D systems) according to the manufacturer’s instructions. Levels of IgM, IgA and IgG_1_ in serum specific for *A. suum* antigen were measured as previously described^45^.

### RNA-sequencing

RNA was extracted from lung and intestinal tissue following homogenization (gentleMACS, Miltenyi Biotech) using miRNAeasy kits (Qiagen) according to the manufacturer’s instructions. RNA was subsequently used for library preparation and 150bp paired-end Illumina NovaSeq6000 sequencing (Novogene, Cambridge, UK). Sequence data was subsequently mapped to the Sus Scrofa (ss11.1) genome and read counts generated which were used to determine DEG using DEseq2. Pathway analysis was conducted using gene-set enrichment analysis (Broad Institute, MA, USA). RNA sequence data from lung and intestinal tissues are deposited at the NCBI Gene Expression Omnibus (Acession numbers: GSE174042 and GSE168840).

### Statistical analysis

All statistical analysis was performed using GraphPad Prism 8, IBM SPSS Statistics 27 or R packages. The data were analyzed using a mixed-model analysis, with diet and infection status as fixed factors and pen and pig as random factors, or with two-way ANOVA and t-tests, as indicated. Where appropriate, time was included as an additional fixed factor to account for repeated measures. One pig (in the *A. suum* + PAC group) was excluded from analysis as it displayed post-mortem pathology indicative of ileitis and aberrant values on several immunological assays. Shapiro-Wilk and Kolmogorov-Smirnov tests were used to tests for assumptions of normality in analyses, and square-root transformations were used to approximate normal distributions when appropriate. For gut microbiota α-diversity analysis, pairwise Wilcoxon Rank Sum Test from the R package was used to obtain Benjamin–Horchberg corrected p-values. Statistical analysis for distance-based redundancy analysis (db-RDA) was done by using permutational ANOVA in the R package vegan. Volcano Plots were created using VolcaNoseR^46^, and principal component analysis was carried out using ClustVis^47^.

## Results

### Proanthocyanidins exert minor effects on systemic antibody and inflammatory biomarkers

Acute infections with *A. suum* induce potent immune reactions in the lungs and intestine before expulsion of the majority of the invading larvae starting from around day 18 p.i.^48^. We fed pigs either a control diet or a diet supplemented with 1% PAC for 14 days, before half the pigs in each group were infected with 5000 embryonated *A. suum* eggs. Mean larval burdens at day 14 p.i. were not altered by PAC supplementation (2914 larvae ± 928 larvae (mean ± SD, *n* = 6) in control-fed pigs and 3155 ± 1057 larvae (mean ± SD, *n* = 5) in PAC fed pigs). In order to examine whether PAC influenced the development of the immune response to *A. suum*, we first examined serological markers of infection in the different treatment groups. *A. suum* infection resulted in a significant increase in serum IgM, IgA, IgG1 specific for *A. suum* antigenic extracts compared to un-infected groups (**Figure 1A-C**). PAC supplementation increased the levels of all three antibody classes in infected pigs, however the effect of diet was not statistically significant. To assess the effect of both *A. suum* and dietary PAC on systemic inflammation, we quantified C-reactive protein (CRP) levels in serum. CRP levels on day 0 p.i. (i.e. after 14 days of PAC supplementation) were significantly lower in PAC fed pigs compared to controls (**Figure 1D**). However, CRP levels measured on day 14 p.i. (i.e. after 28 days of PAC supplementation) were no longer affected by diet, and infection had no impact on CRP levels (**Figure 1E**). Thus, whilst dietary PAC appeared to exert transient anti-inflammatory properties in uninfected pigs, PAC had little capacity to alter systemic antibody production induced by infection.

**Figure 1.**
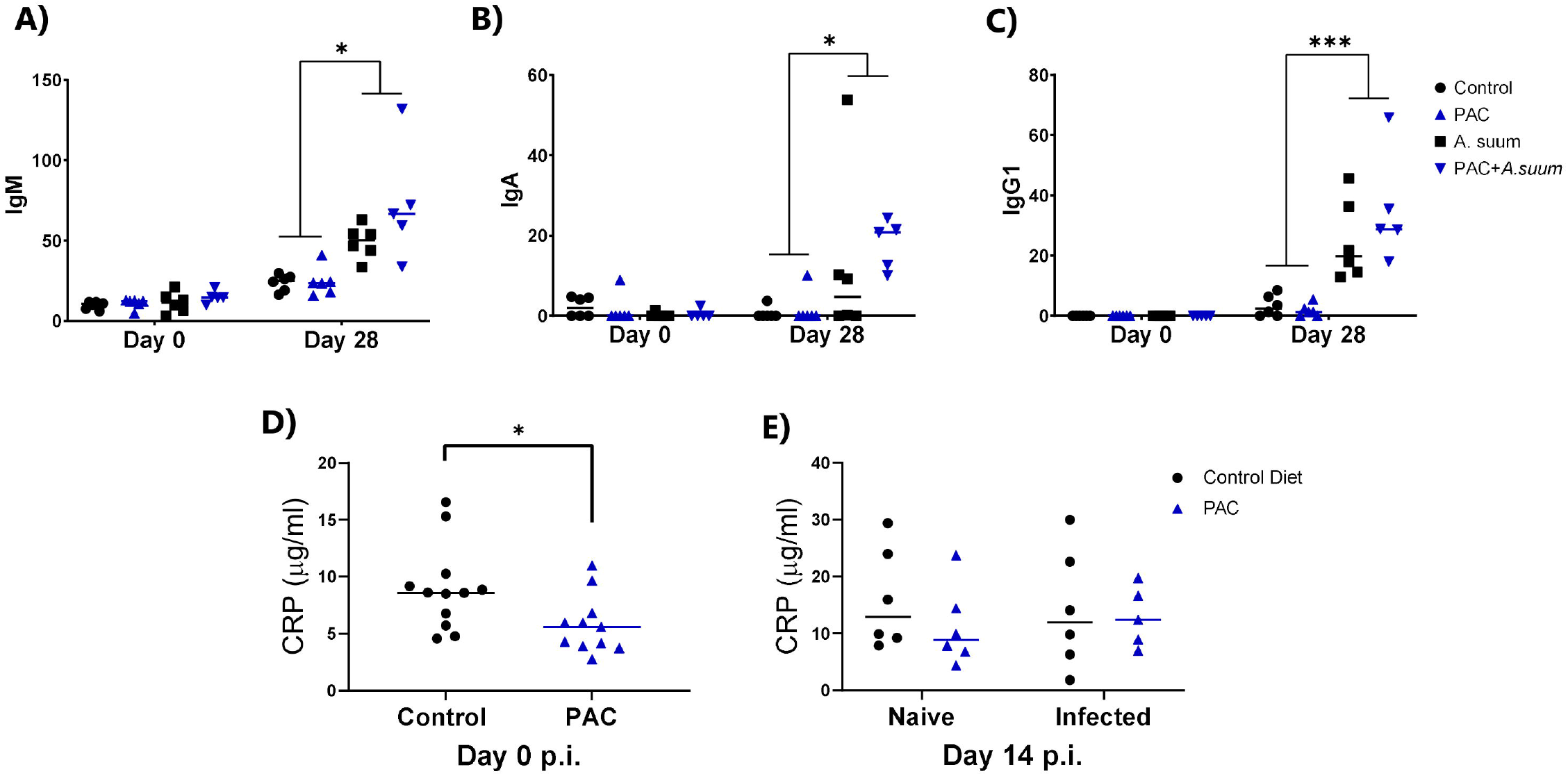
Dietary proanthocyanidins exert limited effects on systemic antibody levels and inflammatory biomarkers. Serum levels of **A)** IgM, **B)** IgG1 and **C)** IgA specific for *Ascaris suum* antigen on day −14 pre-infection (i.e. day of arrival) and day 14 post-infection (p.i.) with *A. suum*. **D)** C-reactive protein (CRP) levels at day 0 p.i. (i.e. after 14 days of proanthocyanidin (PAC) supplementation) and **E)** day 14 p.i. (**Mixed model analysis or t test**, ***p*** < **0.05**, \*\****p*** < **0.01**, \*\*\****p*** < **0.001,** *n* = 6 pigs per group, except *n* = 5 pigs in PAC+*A. suum* group, and n=12 pigs per group in Panel D prior to *A. suum* infection).

### Impact of *Ascaris suum* infection and dietary proanthocyanidins on Th1, Th2 and granulocytic responses in pulmonary and gut tissues

*A. suum* infection induced significant cellular changes in the BAL fluid and tracheal-bronchial lymph nodes (LN). In the LN, the proportion of CD3^+^ T cells was significantly decreased when comparing infected pigs to controls, similar to what has been observed in mice^25^ (**Figure 2A**). The proportions of CD3^+^CD4^+^, CD3^+^CD4^+^T-bet^+^ (Th1) and CD3^+^CD4^+^GATA3^+^ (Th2) T-cells were not significantly different across treatment groups, however Th2/Th1 ratios clearly demonstrated a strong Th2-polarized immune response as a result of *A. suum* infection (**Figure 2B-E**). Moreover, infection markedly induced granulocytosis in BAL fluid, and eosinophilia in jejunal tissues. PAC did not alter granulocyte numbers in BAL, and whilst intestinal eosinophils were numerically higher in infected pigs fed PAC compared to those fed the control diet this difference was not significant (**Figure 2F-G**). Taken together, these findings indicate that *A. suum* induced a strong type-2 biased cellular response in the lungs and intestinal tissues, which was not significantly altered by concurrent PAC consumption.

**Figure 2.**
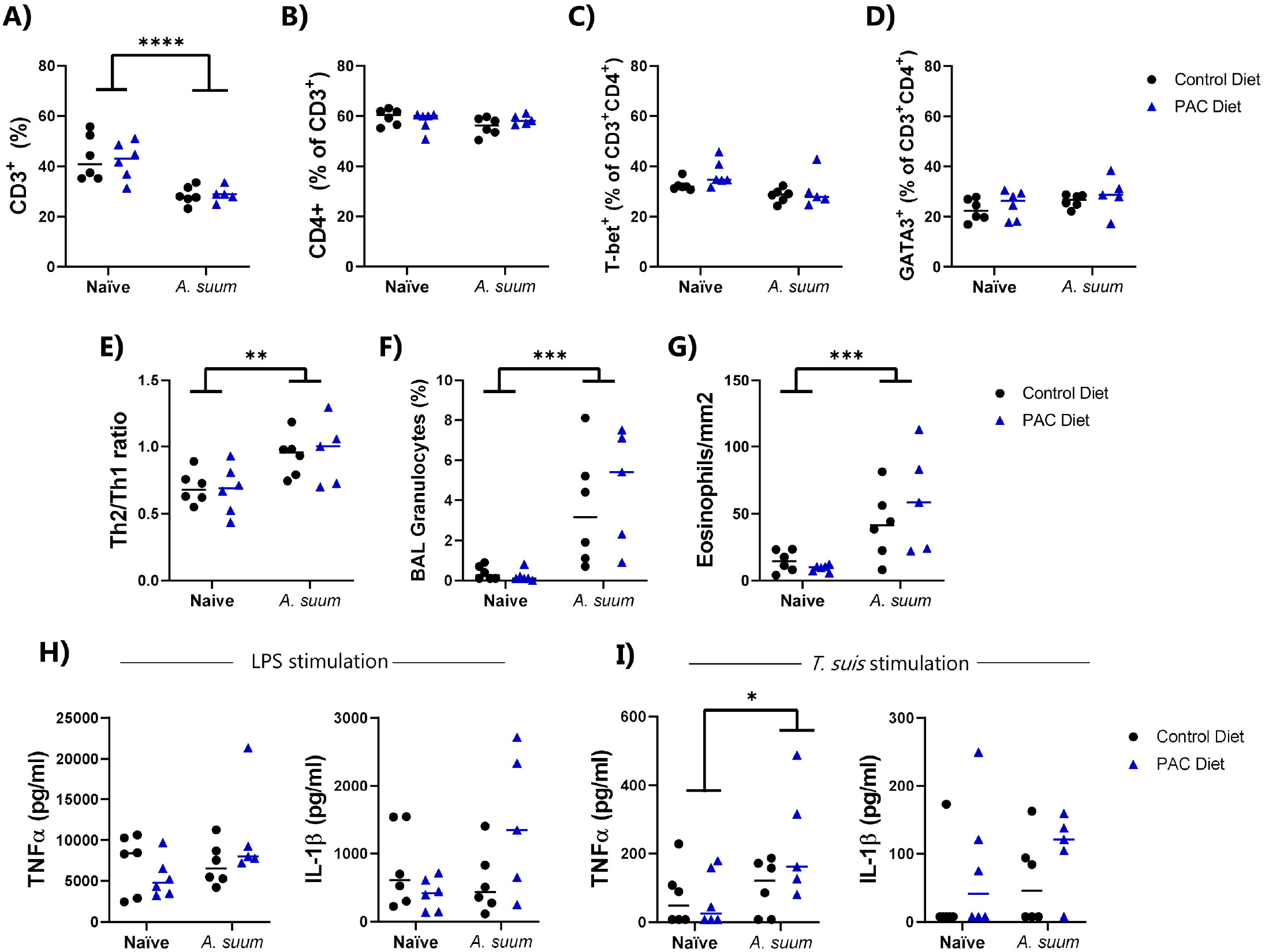
Effects of *Ascaris suum* infection and dietary proanthocyanidins on cellular responses in the lungs and intestine. **A-E)** Proportions of CD3^+^ cells, CD4^+^ T cells, T-bet^+^ Th1 cells, GATA3^+^ Th2 cells, and Th1/Th2 ratios in lung lymph nodes (LN) on day 14 post-infection (p.i.). **F)** Lung granulocytosis in broncho-alveolar lavage (BAL) fluid at day 14 p.i. **G)** Eosinophils in mid-jejunum tissues at day 14 p.i.) TNFα and IL-1β secretion *ex vivo* in alveolar macrophages stimulated with lipopolysaccharide (LPS) (**H)** or *Trichuris suis* antigens **(I).** PAC: proanthocyanidins (**Mixed model analysis,** \****p*** < **0.05**, \*\****p*** < **0.01**, \*\*\****p*** < **0.001**, \*\*\*\****p*** < **0.0001**, *n* = 5-6 pigs per group).

### Concomitant *Ascaris suum* infection and dietary proanthocyanidins tend to enhance cytokine secretion in alveolar macrophages stimulated *ex vivo*

To further assess the effects of infection and dietary PAC on the profile of lung immune cells, alveolar macrophages were isolated from BAL of all pigs in each treatment group and stimulated *ex vivo*. Cells were first stimulated with LPS to assess how diet and infection may influence secretion of the pro-inflammatory cytokines TNFα and IL-1β. Infection status did not significantly influence LPS-induced secretion of these cytokines. Whilst levels of LPS-induced TNFα and IL-1β were lower in uninfected pigs fed PAC, the secretion of these cytokines tended to be higher in infected pigs fed PAC, albeit not significantly so (*p* = 0.15 for interaction between diet and infection status; **Figure 2H**). To explore if *ex vivo* inflammatory responses to parasite antigens were modulated by PAC or infection, macrophages were first stimulated with *A. suum* E/S products, however no cytokine secretion was observed (data not shown). We therefore also stimulated cells with E/S from another porcine helminth, *T. suis*, which we have previously shown to be a stronger activator of innate cytokine production^49^. Interestingly, alveolar macrophages isolated from infected pigs secreted significantly higher levels of TNFα when activated with *T. suis* E/S products (**Figure 2I**). Similarly to LPS-stimulated cells, macrophages isolated from infected pigs fed PAC secreted higher levels of TNFα and IL-1β compared to macrophages isolated from any other treatment groups, albeit not significantly (**Figure 2I**). These findings suggest that *A. suum* infection primed macrophages to be more responsive to stimulation from heterologous parasite antigens (but not LPS), whilst PAC has only minor systemic immuno-stimulatory effects on cytokine production during infection.

### Transcriptional profiling of gut and lung tissues reveals modulatory effects of proanthocyanidins during *Ascaris suum* infection

#### Jejunum transcriptional responses

To explore in more detail if PAC may influence the immunological response to *A. suum* infection, we conducted RNA-sequencing of jejunal and lung tissues. Both *A. suum* and PAC treatment strongly modulated gene expression in the intestine as compared to controls (**Figure 3 and 4**). Principal component analysis showed a clear clustering of biological replicates according to infection status (**Figure 3A**). *A. suum* infection significantly downregulated the expression of genes such as the aldehyde dehydrogenase-encoding *ALDH1B1*, and the sodium-channel encoding *SCN8A*. Interestingly, three of the top-ten downregulated genes as a result of infection were related to circadian rhythm (*PER3, PER2, NOCT*) (**Figure 3B-C**). Moreover, *A. suum* infection significantly upregulated the expression of interleukins *IL4, IL9, IL10, IL21*, the eosinophil marker *EPX*, as well as TCR related genes *CD28* and *CD80* in intestinal tissue. Furthermore, we noted strong upregulation of genes involved in aryl hydrocarbon receptor (AHR)-signaling including *ARNTL* as well as smooth muscle contraction (*P2RX1*), which may relate to the increased intestinal motility observed during the immune reaction to *A. suum* larvae^50^ (**Figure 3B-C**). Analysis of gene pathways modulated by infection revealed that pathways related to peroxisome function, as well as the metabolism of fatty acids and glycerolipids were significantly suppressed, suggesting a profound modulation of nutrient metabolism due to larval colonization of the intestine (**Figure 3D**). Unsurprisingly, the main up-regulated gene pathways were related to immune function, such as the IL-2, IL-4, and T-cell receptor related pathways, as well as granulocyte and B-cell signaling (**Figure 3D**). Thus, consistent with the pulmonary and intestinal eosinophilia, *A. suum* induced a type-2 inflammatory reaction concomitant with pathophysiological responses related to the changed mucosal environment induced by larval antigens.

**Figure 3.**
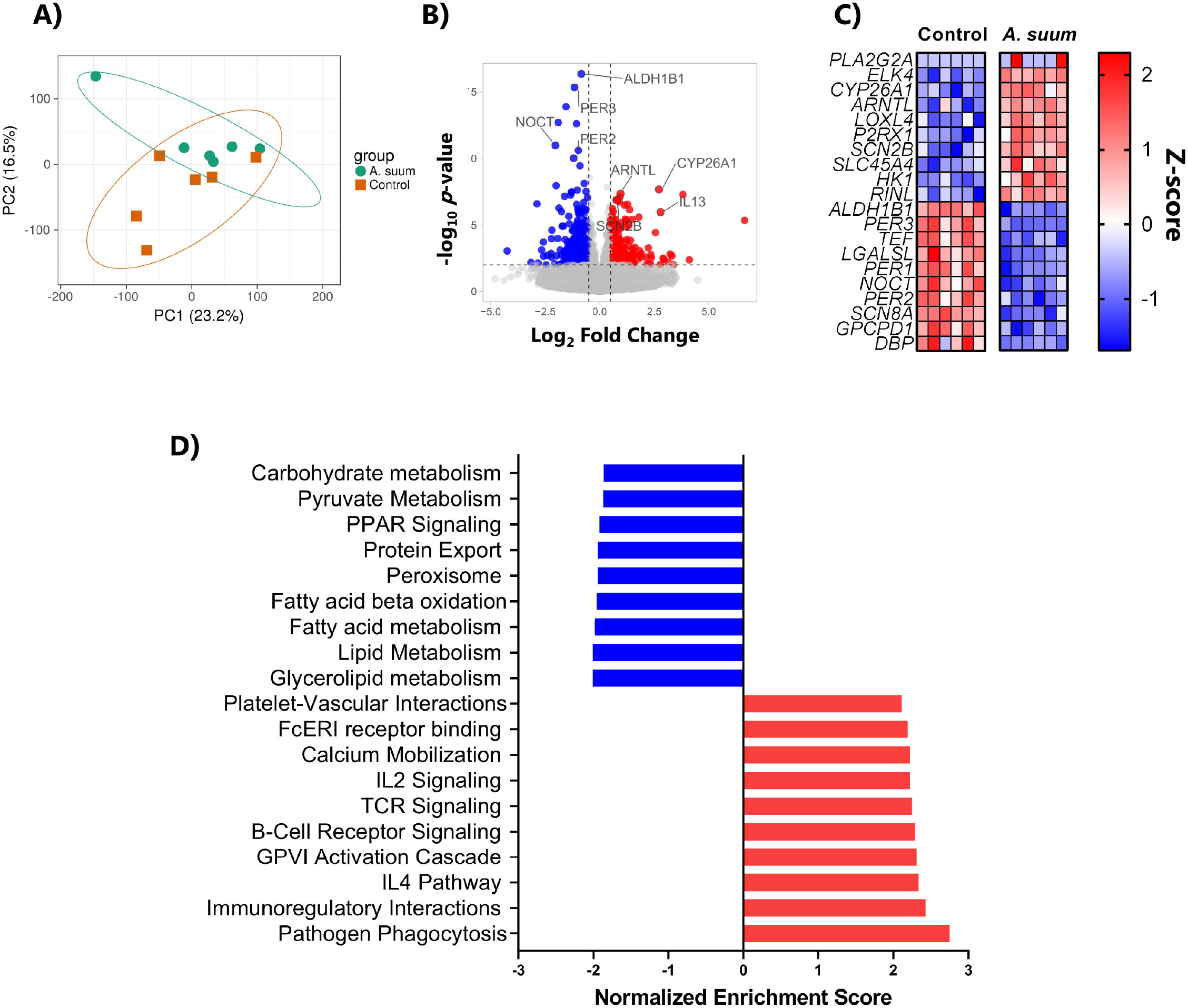
Modulation of gene expression and transcriptional pathways in intestinal tissue by *Ascaris suum* infection. **A)** Clustering of *A.suum*-infected and control groups as demonstrated by principal component analysis. **B)** Volcano plot showing differentially expressed genes resulting from *A. suum* infection **C)** Top ten up- and down-regulated genes identified as a result of *A. suum* infection. (*n* = 6 pigs per group). **D)** Significantly up- and down-regulated pathways (*p* < 0.01; Q < 0.1) identified by gene-set enrichment analysis as a result of *A. suum* infection.

**Figure 4.**
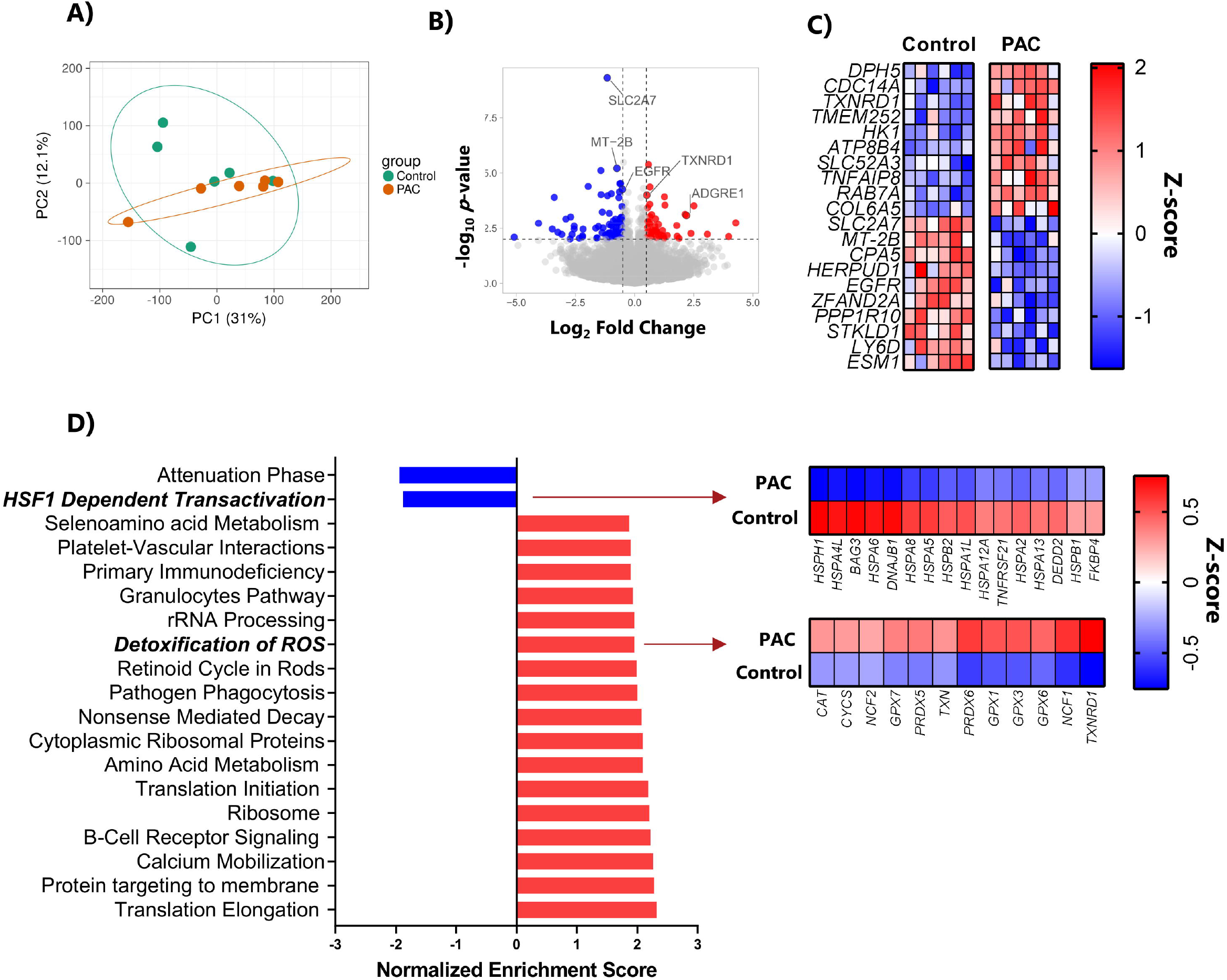
Modulation of gene expression and transcriptional pathways in intestinal tissue by dietary proanthocyanidins. **A)** Clustering of the two dietary groups as demonstrated by principal component analysis in uninfected pigs. **B)** Volcano plot showing differentially expressed genes resulting from dietary proanthocyanidin (PAC) supplementation. **C)** Top ten up- and down-regulated genes identified as a result of dietary PAC supplementation in uninfected pigs. (*n* = 6 pigs per group). **D)** Significantly up- and down-regulated pathways (*p* < 0.01; Q < 0.1) identified by gene-set enrichment analysis (GSEA) as a result of dietary PAC supplementation. Highlighted are the HSF1 Dependent Transactivation and Detoxification of ROS pathways, showing enriched genes as identified by GSEA analysis.

In uninfected pigs, dietary PAC resulted in a distinct clustering of treatment groups based on diet as assessed by principal component analysis (**Figure 4A**). Downregulated genes included the glucose transporter *SLC2A7*, which has previously been shown to be inhibited by polyphenols in cellular models^51^, as well as *MT-2B*, encoding a metallothionein protein known to be associated with intestinal inflammation and oxidative stress in mice^52^. Interestingly, we also noted downregulation of *EGFR*, encoding the epidermal growth factor receptor (**Figure 4B-C**). Significantly upregulated genes included *TXNRD1*, encoding thioredoxin reductase 1, a protein involved in suppression of reactive oxygen species (ROS), as well as genes involved in cellular endocytic processes (*RAB7A*) and extracellular matrix remodeling (*COL6A5*) (**Figure 4C**). Consistent with this, gene pathways related to metabolic processes, such as translation elongation and ribosome function, were significantly enriched (**Figure 4D**). Interestingly, we noted that a number of pathways that were related to immune function (and that were also induced by *A. suum)*, were upregulated by PAC. These included pathways related to granulocyte function and B-cell signaling, indicative of an immune-stimulatory effect of PAC. Furthermore, we observed a significant upregulation of pathways related to both detoxification of ROS and selenoamino acid metabolism, suggestive of enhanced antioxidant responses in PAC-fed pigs. Notably, we also observed a strong downregulation of pathways related to heat shock responses, which are normally induced by cellular stressors and offer protection against tissue injury^53^ (**Figure 4D**). Collectively, these data suggest that dietary PAC have significant effects on intestinal metabolism and function as a cytoprotective agent in the intestinal mucosa, by inducing antioxidant responses and regulating responses to cellular stressors.

Given that PAC appeared to induce transcriptional pathways with functions in immunity and inflammation, we next asked whether concurrent PAC consumption could modulate the intestinal transcriptomic response to *A. suum* infection. We observed that within infected pigs, there was once again a clear clustering according to diet based on principal component analysis (**Figure 5A**). Inspection of genes differentially expressed in infected, PAC-fed pigs, relative to infected pigs fed the control diet, revealed that expression of genes involved in intestinal nutrient metabolism were increased, such as *ORAI2* and *AGTR1*, which both play a role in calcium uptake^54, 55^. Consistent with the suppression of *EGRF* expression in uninfected pigs fed PAC, the expression of a number of genes related to EGF signaling, including *BTC* and *AREG*, were downregulated. *AREG* encodes amphiregulin, a cytokine involved in type-2 inflammation induced by a number of different helminth species^56^ (**Figure 5B)**. In agreement with the data showing an enrichment of antioxidant pathways in uninfected pigs fed PAC, we noted that PAC supplementation during infection also resulted in the upregulation of the oxidative stress pathway, which included significant enrichment of *SOD3, GPX3* and *NQO1* - genes encoding proteins with known anti-oxidant properties (**Figure 5C**). Down-regulated pathways in infected pigs fed PAC were mainly related to metabolic activity such as cholesterol metabolism, but these were not significant following FDR adjustment (data not shown). Collectively, these data show that PAC exert a significant influence on the intestinal transcriptional environment during enteric helminth infection mainly by promoting the transcription of genes involved in regulating oxidative stress and nutrient metabolism.

**Figure 5.**
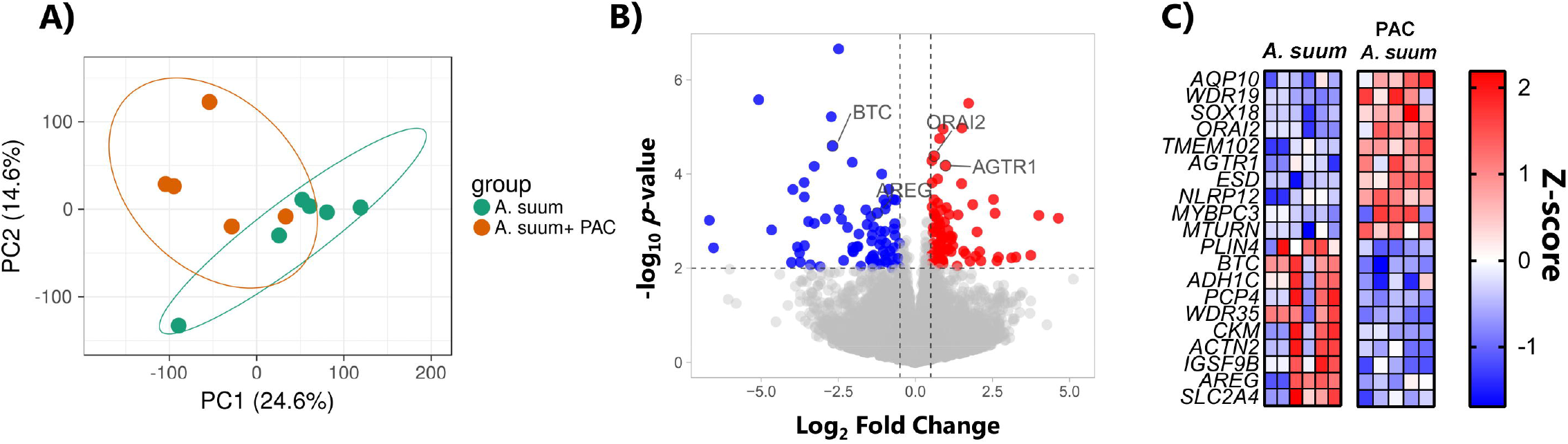
Modulation of gene expression and transcriptional pathways in intestinal tissue by dietary proanthocyanidin supplementation in *Ascaris suum*-infected pigs. **A)** Clustering of the dietary groups within *Ascaris suum*-infected pigs as a result of dietary proanthocyanidins (PAC) as demonstrated by principal component analysis. **B)** Volcano plot showing differentially expressed genes resulting from dietary PAC supplementation in *A. suum-infected* pigs. **C)** Top ten up- and down-regulated genes identified as a result of dietary PAC supplementation in *A. suum*-infected pigs. (*n* = 6 pigs in *A. suum* group, *n* = 5 pigs in PAC+*A. suum* group).

#### Lung transcriptional responses

Next, transcriptional profiling of the lungs by RNA-sequencing was performed to investigate the effect of larval migration in the lungs and the potential impact of dietary PAC on gut-lung interplay. In comparison to the intestine, the modulation of gene expression in the lungs was only modestly modulated by both *A. suum* infection and/or PAC supplementation. Interestingly, as was the case in the jejunum, *A. suum* infection regulated the expression of numerous genes related to circadian rhythm. Notably, *PER1, PER2, PER3, NR1D1, NR1D2* and *DBP* were suppressed, whereas *NPAS2, ARNTL* were significantly upregulated (**Figure 6A**). A number of studies have touched upon the importance and complex interplay between circadian rhythm, immune regulation and parasite-host interactions^57^. Of note, *ARNTL* was also significantly upregulated by PAC (**Figure 6B**). In coherence with the above described granulocytosis in the lungs in BAL fluid, *A. suum* infection upregulated the expression of *CCR3*, which is essential for eosinophil recruitment. Infected pigs fed PAC had significantly higher expression levels of genes related to innate immune function (*CD209* and *OAS2)*, and connective tissue growth factor (*CTGF*) in lung tissues compared to infected pig fed a control diet (**Figure 6C**). CTGF is involved in wound repair and tissue healing, suggesting a protective effect of PAC during *A. suum* infection. Intriguingly, the expression of the oxidative stress inducer *ALOX15* was significantly increased by *A. suum* infection, but was significantly down-regulated in infected pigs fed a PAC diet, which supports previously described reports of PAC acting as a lipoxygenase inhibitor^58^ (**Figure 6A and C**). Thus, *A. suum* infection induced marked transcriptional responses in the lungs but somewhat less than compared to intestinal tissues, which may indicate that lung homeostasis is somewhat restored by day 14 p.i. when the migrating larvae have returned to the intestine. Furthermore, dietary PAC induced smaller transcriptional changes in the lung compared to the intestine but may ameliorate wound healing and antioxidant status during infection.

**Figure 6.**
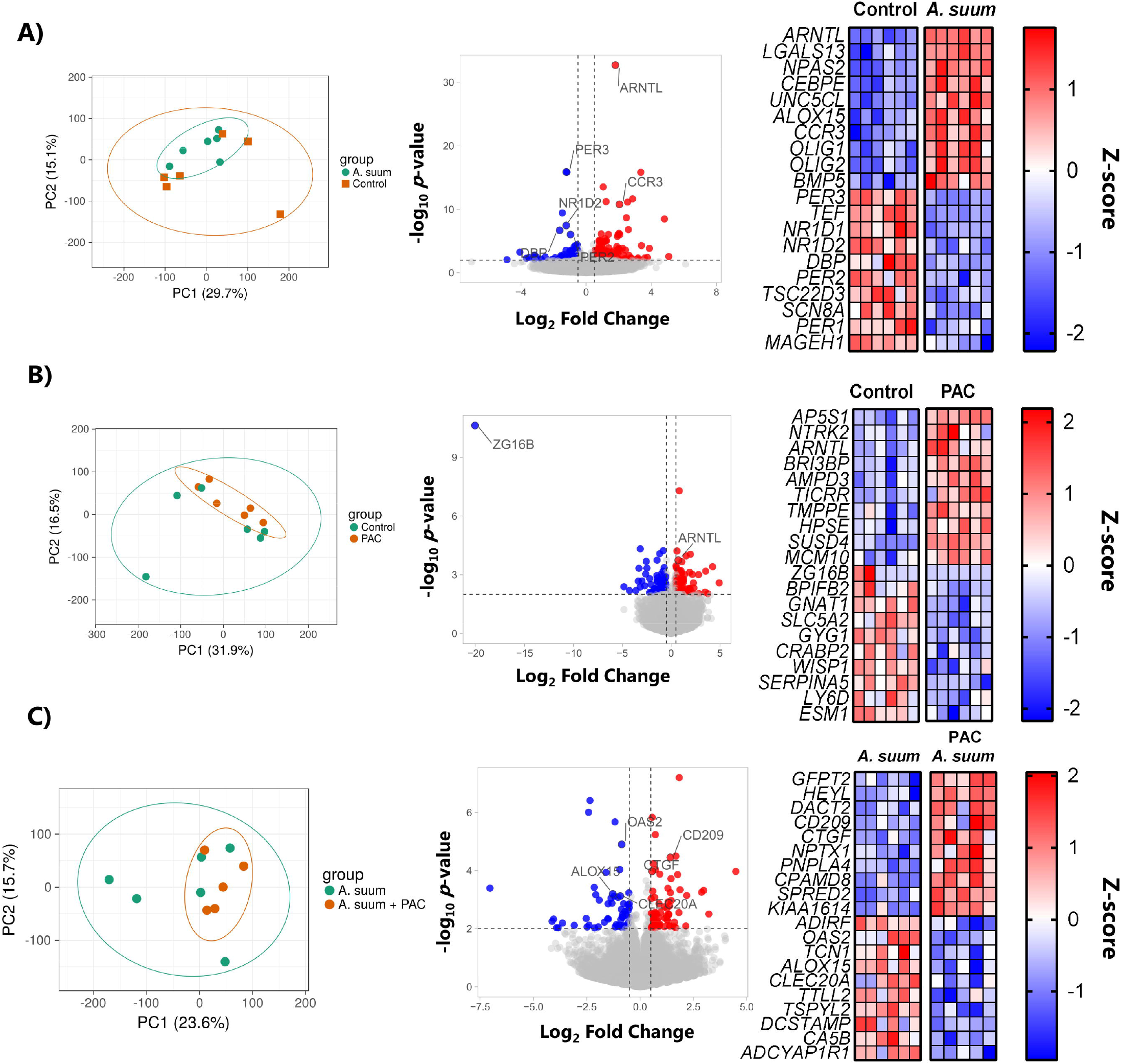
Modulation of gene expression in lung tissue by *Ascaris suum* infection and dietary proanthocyanidins. Effects on lung gene expression as shown by principal component analysis, volcano plot of differentially expressed genes and top ten up- and down-regulated genes identified as a result of **A)** *Ascaris suum* infection in pigs fed the control diet, **B)** dietary proanthocyanidins (PAC) in naïve pigs and **C)** dietary PAC in *A. suum*-infected pigs. (*n* = 6 pigs per group, except *n* = 5 pigs in PAC+*A. suum* group).

### *Ascaris suum* infection and proanthocyanidins alter gut microbiota composition with limited effect on short chain fatty acids

Previous studies have indicated that immunomodulatory and anti-inflammatory effects of PAC may derive from changes in the GM and associated metabolite production^17^. Furthermore, *A. suum* and other helminths can markedly change host GM composition^59^. Therefore, to explore whether the observed transcriptomic changes induced by diet and infection were accompanied by GM changes, we used 16S rRNA gene amplicon sequencing to characterize both the small and large intestinal GM composition. We initially analyzed the GM composition in the jejunum, at the main site of *Ascaris* infection. Neither *A. suum* nor dietary PAC altered α-diversity (data not shown). Changes in β-diversity were apparent primarily as a result of *A. suum* infection (*p* < 0.05 by distance-based redundancy analysis; **Figure 7A**), with differential abundance analysis on genus level indicating an enrichment in *Lactobacillus* spp. in infected pigs (**Figure 7B**). Moreover, *A. suum* infection decreased the abundance of *Facklamia* spp. (*p* = 0.053 by mixed model analysis; **Figure 7C**). In contrast, PAC did not have a significant effect on the small intestinal GM, with no changes in β-diversity between PAC-fed pigs and control pigs (*p* > 0.05 by distance-based redundancy analysis; **Figure 7A**). However, we did note that, within *A. suum-*infected pigs, those animals fed PAC tended to have a higher abundance of amplicon sequences corresponding to *Limosilactobacillus reuteri* (**Figure 7D**). *L. reuteri* has been associated with beneficial probiotic and anti-inflammatory effects, and plays a role in the prevention of microbial translocation and inhibits colonization of pathogenic bacteria^60, 61^.

**Figure 7.**
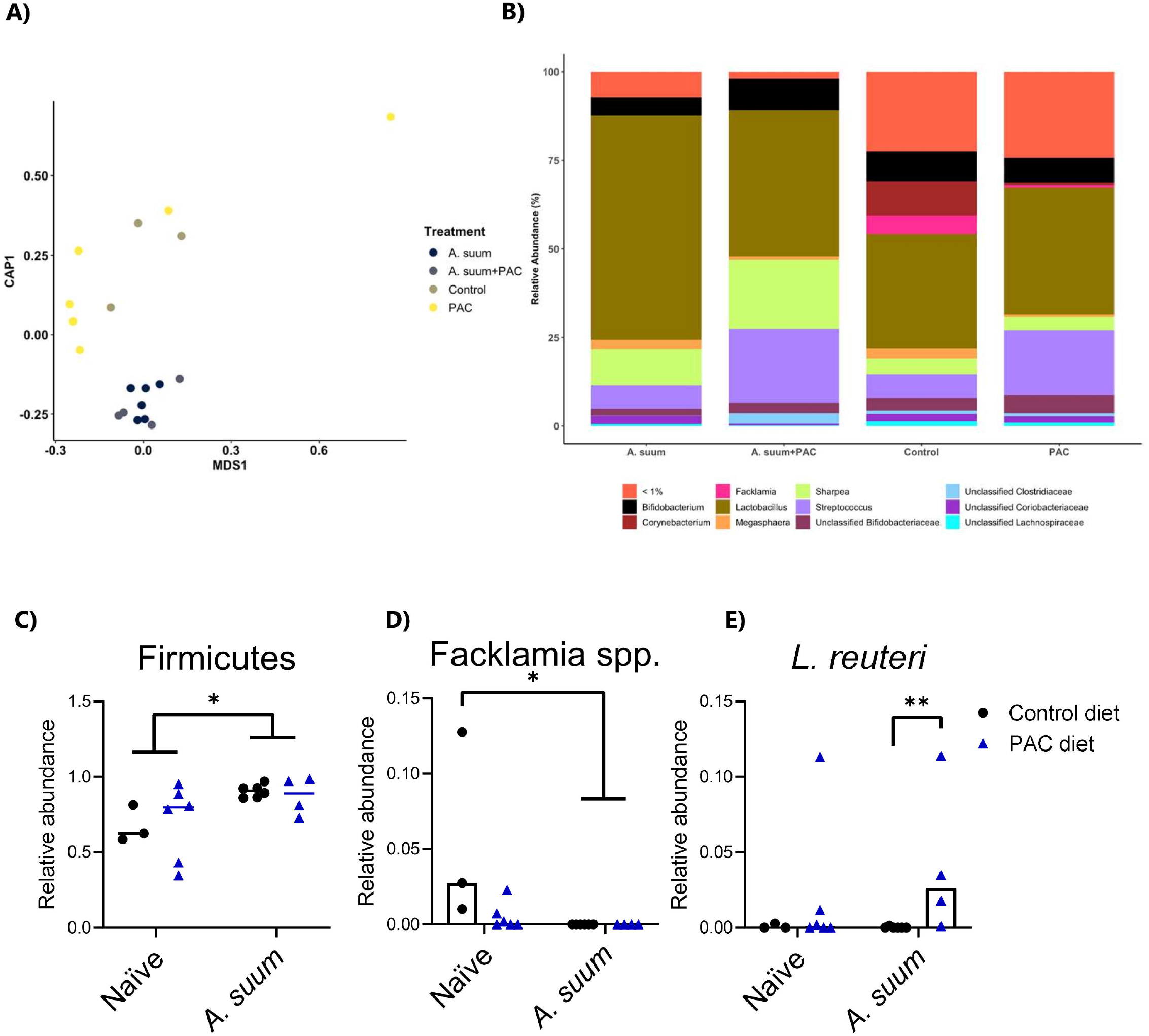
Changes in gut microbiota composition in the small intestine due to dietary proanthocyanidins and *Ascaris suum* infection. **A)** Changes in β-diversity in the small intestine, as identified by distance-based redundancy analysis where a significant effect of *Ascaris suum* compared to all other treatment groups was identified. No effect on β-diversity was reported when comparing PAC to control-fed naïve pigs **B)** Relative abundance at genus level in naive or *A. suum*-infected pigs fed a control diet or PAC-supplemented diet. Relative abundance of **C)** *Facklamia* spp. and **D)** *Limosilactobacillus reuteri* in naive or *A. suum*-infected pigs fed a control diet or PAC-supplemented diet, as identified by differential abundance analysis and mixed-model analysis. (*n* = 3 pigs in control group, *n* = 6 pigs in *A. suum* group, *n* = 6 pigs in PAC group, and *n* = 5 pigs in PAC+*A. suum* group).

In the colon, we found that PAC had the largest effect on the GM composition, consistent with the notion that PAC are extensively metabolized by, and can modulate, the large intestine microbiome (*p* < 0.05 for β-diversity comparison between PAC and control group by distance-based redundancy analysis; **Figure 8A-B**). PAC tended to decrease the abundance of *Bifidobacterium* in both naïve and infected pigs (**Figure 8C**). Notably, the abundance of sequences closely related to *Bifidobacterium thermacidophilum* was significantly increased by *A. suum*, but concomitant PAC supplementation significantly suppressed this effect (**Figure 8D**). The reduction of *Bifidobacterium* in pigs fed PAC contrasts to a previous study in pigs showing that PAC increased the growth of this taxa^15^. However, similar to the trend in the small intestine, PAC supplementation resulted in the significant increase of *L. reuteri* abundance in the colon of both naïve and infected pigs (**Figure 8E**). Interestingly, *A. suum* infection increased the abundance of *Lactobacillus* spp. in the colon whilst significantly decreasing the abundance of *Turicibacter* spp. (**Figure 8F and G**). However, β-diversity was not different between *A. suum* and control groups in colon, indicating that the effects of infection on GM composition were mostly limited to the predilection site (the small intestine). Finally, we investigated if the colonic GM changes were accompanied by changes in the concentrations of SCFA in the distal colon. Neither PAC nor *A. suum* infection altered levels of acetic acid, propionic acid, n-butyric acid or D-lactic acid (**Supplementary Figure 3**). However, we observed that dietary PAC decreased the concentrations of the branched-chain fatty acids iso-valeric acid (*p* < 0.05) and iso-butyric acid (*p* = 0.0616), which may relate to altered protein metabolism or colonic transit time^62^, and is consistent with our previous work on pigs fed a polyphenol-enriched diet^63^ (**Figure S1**). Taken together, these results indicate distinct effects of *A. suum* infection and PAC on specific bacteria taxa in a site-specific manner.

**Figure 8.**
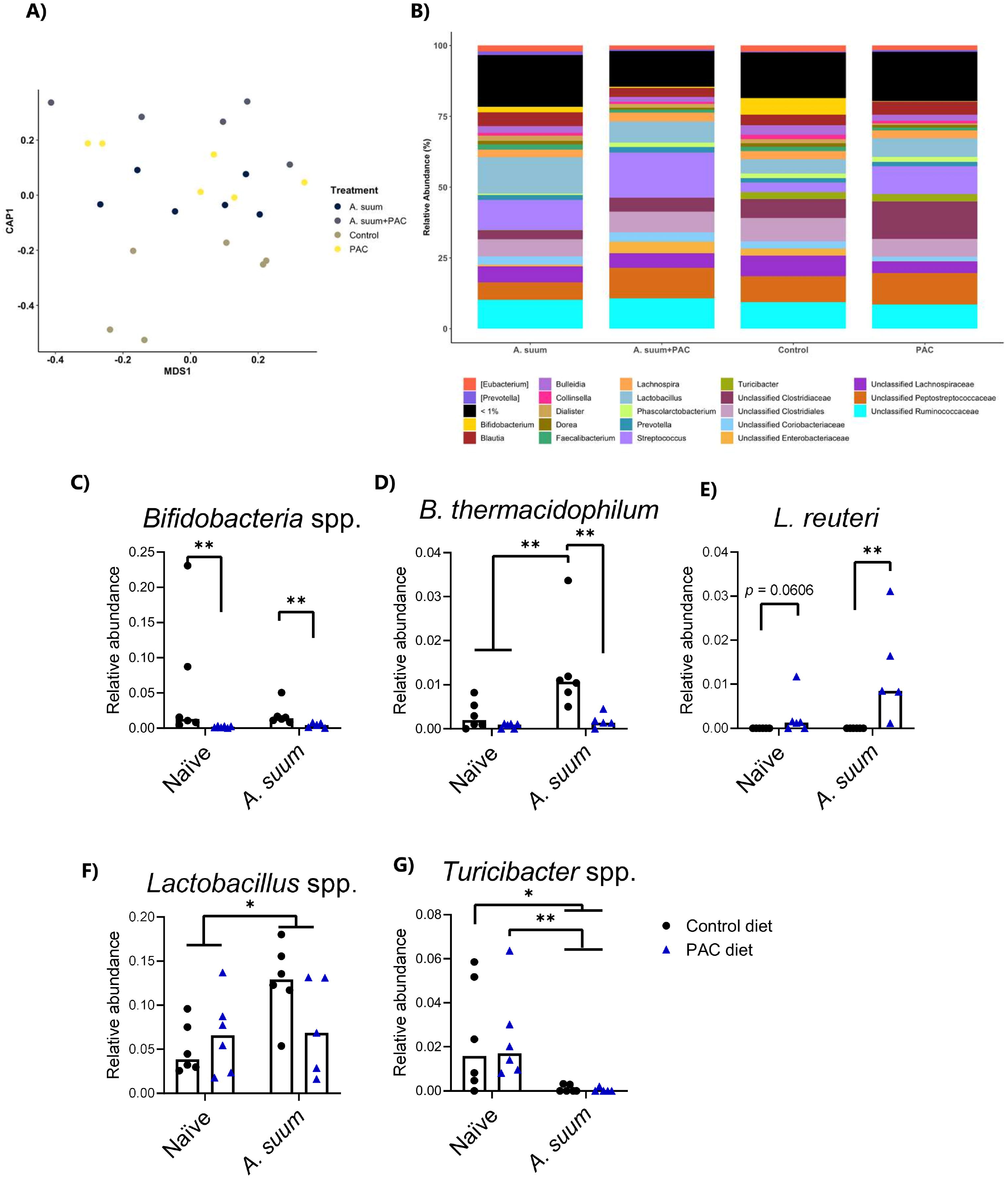
Changes in gut microbiota composition in the proximal colon due to dietary proanthocyanidins and *Ascaris suum* infection. **A)** Changes in β-diversity in the proximal colon as identified by distance based redundancy analysis, where an effect of proanthocyanidins (PAC) compared to control was identified.. **B)** Relative abundance at genus level in naive or *Ascaris suum*-infected pigs fed a control diet or PAC-supplemented diet. Relative abundance of **C)** *Bifidobacteria* spp., **D)** *B. thermacidophilum*, **E)** *Limosilactobacillus reuteri*, **F)** *Lactobacillus* spp. and **G)** *Turicibacter* spp. in naive or *A. suum*-infected pigs fed a control diet or PAC-supplemented diet, as identified by differential abundance analysis and mixed-model analysis (\****p*** < **0.05**). (*n* = 6 pigs per group, except *n* = 5 pigs in PAC+*A. suum* group).

## Discussion

The immuno-modulatory effects of PAC have been investigated in numerous studies but their mode of action and impact on immune function is still not fully understood. Furthermore, only limited knowledge has been attained on the effects of PAC on type-2 immune response, which plays a central role during helminth infections and may be relevant for inflammatory disorders, such as food allergies and ulcerative colitis. Therefore, we used here a model of *A. suum* infection in pigs, which offers a unique opportunity to explore the modulation of parasite-induced inflammation in multiple tissues by dietary components.

Initial assessment of the systemic effects of PAC and *Ascaris* infection were demonstrated by monitoring serum antibody levels and the acute-phase protein CRP, a marker for systemic inflammation. *A. suum* infection resulted in a significant increase in serum antibodies, which were further enhanced by dietary PAC, albeit not significantly. Interestingly, significantly lower CRP levels were observed after 14 days of PAC supplementation, although this effect subsided by the end of the study. Thus, PAC had limited effects on parasite-induced antibody levels, and prolonged PAC supplementation did not appear to persistently alter inflammatory markers in serum.

The gut-lung axis is gaining increasing interest in numerous research fields, and the migratory characteristics of *A. suum* render the investigation of gut-lung interplay greatly relevant in this model. Here, we showed that *A. suum* infection induced granulocytosis in the lungs, and a Th2 polarized immune response was clearly demonstrated by Th2/Th1 T-cell ratios. PAC and *A. suum* in isolation, upregulated a number of similar genes, notably genes related to the circadian rhythm, such as *ARNTL*. Interestingly, a recent study showed that *ARNTL*, also known as *BMLA1*, was fundamental for the time-of-the-day dependent expulsion efficiency of *Trichuris muris* infection in mice^64^. Furthermore, a study conducted in pigs also demonstrated an association between *ARNTL* and adult worm burden^65^. However, in contrast to murine studies, which have demonstrated a role for PAC in suppressing allergic responses in the lungs, we did not find a modulatory effect of PAC on the type-2 cellular response to *A. suum* infection. *Ex vivo* stimulation of lung macrophages by LPS or helminth antigens indicated a tendency of higher cytokine secretion levels in macrophages isolated from infected pigs fed PAC. Moreover, although we observed transcriptional changes in the lungs of infected pigs that are reflective of type-2 inflammation, these did not appear to be markedly altered by concurrent PAC intake. The exception was an indication of regulation of several genes such as *CTGF*, and *ALOX15*, which could suggest that PAC may augment wound-healing and anti-oxidant status in lung tissues during *A. suum* infection. Thus, in our model, dietary PAC had limited capacity to regulate lung immune function during helminth infection, although further studies to elucidate whether PAC may potentiate protection towards secondary airway infection during *A. suum* infection may be relevant.

We next assessed the impact of *A. suum* infection and PAC at the predilection site of infection, the small intestine. *A. suum* infection induced stereotypical intestinal eosinophilia, which was equivalent in both dietary groups. We had previously shown that eosinophilia in the jejunum of *A. suum-*infected pigs could be potentiated by a polyphenol-enriched diet containing 5 % grape pomace^63^. Grape pomace may contain several phytonutrients and fibrous components such as lignin, which could contribute to synergistic effects, whereas the PAC diet in the present study was composed only of purified PAC oligomers from grape seed extract. This may explain the discrepancy between these results. Transcriptomic analysis of intestinal tissues revealed that a number of genes and pathways were regulated by both infection and PAC supplementation. As expected, *A. suum* induced the upregulation of type 2 immune related genes and pathways, as well as having an important impact on nutrient metabolism-related genes. Notably, PAC and *A. suum* in isolation were both able to modulate transcriptional pathways related to immune function and antioxidant activity. Interestingly, PAC increased the expression of protein-encoding genes with cytoprotective functions against oxidative stress, suggesting a role in improving gut heath by minimizing cellular stress during inflammation. The antioxidant effect of PAC could be caused by the absorption of PAC-derived metabolites, produced as a result of microbial metabolism. Although PAC are known to remain relatively stable until they reach the large intestines, PAC molecules with low mDP may also be absorbed in the small intestines^66, 67^. Furthermore, PAC and their metabolites may exert direct interactions with the gut mucosa and thus epithelial cells, as described in numerous cell-based studies. PAC may intervene as scavengers of free radicals due to the hydroxyl groups present in their molecular structures, which can neutralize free radicals via electron delocalization^5^. Another mechanism of the protective effects of PAC, may be via the induction of cellular antioxidant defenses by modulating Nuclear factor erythroid 2-related factor 2 (Nrf2)-related genes, which play an important role in regulating cellular resistance to oxidants, such as ROS^68^.

The localized effect of PAC and infection in the intestines was also demonstrated by their impact on the GM. *A. suum* infection caused substantial changes in the GM composition, most notably in the small intestine. This is the first report of alterations in the GM by *Ascaris* in the predilection site of the jejunum, and we found a significantly decreased abundance of *Facklamia* spp. Furthermore, we noted that *A. suum* increased the abundance of lactobacilli in the colon. Consistent with this, an increased abundance of lactobacilli has also been associated with *Heligmosomoides polygyrus* infection in mice^69, 70^. This may potentially result from the increased mucus secretion that is a stereotypical feature of helminth infections which may provide a niche environment for lactobacilli to thrive^71^. Interestingly, the abundance of *L. reuteri* was significantly increased by PAC supplementation in both naïve and infected pigs, suggesting a prebiotic effect, which may have functional implications, given the known role of *L. reuteri* in modifying inflammation. However, PAC also significantly decreased the abundance of *Bifidobacterium* spp., including *B. thermacidophilum*, suggesting a complex regulation of the GM. Notably, the suppressive effect of PAC on *Bifidobacterium* spp. stands in contrast to a previous study showing the opposite effect in pigs fed PAC derived from cocoa^15^. These apparently contradictory findings may potentially be explained by the differing molecular structures of PAC derived from different sources, as well as potential interactions with differing basal diets. Given that PAC appeared to change the GM composition, a key question is whether the immunomodulatory effects of PAC in the intestine derive from direct interactions with PAC and mucosal immune cells during intestinal transit, or whether PAC-derived microbial metabolites are absorbed and exert systemic bioactivity, as has been proposed in previous studies^11, 15^. Given that PAC-related transcriptional changes we observed were localized mainly to the gut, and not the lung, this may support a hypothesis that the activities were derived from direct interactions between PAC and cells at the level of the gut mucosa, consistent with a lack of an effect of PAC on SCFA levels. However, further studies are clearly needed to unravel these mechanistic aspects.

In conclusion, pigs infected with *A. suum* offered a robust model to study the effect of PAC on pathogens that induce a strong, type-2 biased mucosal immune response in pulmonary and intestinal tissues. Both *A. suum* infection and PAC in isolation had similar immunomodulatory capacity, notably by modulating gene pathways related to B-cell function. PAC also affected transcriptional pathways related to oxidative stress by significantly increasing the expression levels of protein-encoding genes with cytoprotective properties. However, the canonical markers of type-2 inflammation, such as eosinophilia and Th2 T-helper cells in the lungs, were not modulated by PAC intake. The limited effects of dietary PAC observed in the lungs is in coherence with a previous study demonstrating no effect of PAC on gene expression levels of various immune-related genes in alveolar macrophages and tracheobronchial lymph nodes isolated from *A. suum* infected pigs, which were dosed with PAC derived from cocoa^29^. Thus, in contrast to some murine studies suggesting beneficial effects of dietary PAC on asthma, our results suggest a restricted ability of PAC to influence the development of Th2 responses in the respiratory tract in pigs. However, the significant modulatory effects of PAC on porcine intestinal gene expression suggest a primarily gut-localized effect of PAC. Thus, PAC may play a role in maintaining gut health during enteric infection in pigs and humans, and further studies to address the functional implications of this diet-infection interaction are highly warranted.

## Acknowledgements

The authors would like to thank Mette Marie Schjelde for excellent laboratory assistance and Charlotte Smith Bonde, Lise-Lotte Christiansen, Penille Jensen, Stine Nielsen and Pankaj Arora for assistance with the animal study.

## Ethical statement

All experiments involving animals were conducted in agreement with the Danish legislation and the Danish Animal Experiments Inspectorate with the license number 2015-15-0201-0076.

