## Supplementary Information for "Dietary Proanthocyanidins Exert Localized Immunomodulatory Effects in Porcine Pulmonary and Gastrointestinal Tissues during *Ascaris suum*-induced Type 2 inflammation"

Contents:

Table S1-S2

Figure S1-S3

**Table S1 Composition of basal diet**

| **Composition of basal feed diet** |
|---|
| **Analytical constituents:**<br>16,3 % Crude protein<br>3,8 % Crude fat<br>3,7 % Lignin<br>5,2 % Raw ash<br>0,99 % Lysine<br>0,29 % Methionine<br>0,81 % Calcium<br>0,49 % Phosphor<br>0,17 % Sodium |
| **Composition:**<br>Wheat (57,85%); Soybean meal (20,84%); Barley (15.00%); Molasses (1.50%); Fatty acid distillates from physical refining, palm (1.50%); Calcium carbonate, chalk (1.47%); Monocalcium phosphate (0.70%); Sodium chloride, feed salt (0.37%); L-Lysine monohydrochloride (3.2.3) (0.29%); Premix Vit. Slagt (0.20%); Methionine 40 (3.1.1) (0.17%); L-Threonine (3.3.1) (0.13%); |
| **Additives, added per kg:**<br>Nutritional:<br>4200 i.e Vitamin A (3a672a)<br>420 i.e. Vitamin D3 (3a671)<br>84 i.e. E-Vit 3a700 all-rac-alpha-tocopheryl acetate<br>84 mg Fe, iron II sulphate (3b103)<br>15 mg Cu, copper II sulphate (3b405)<br>42 mg Mn, manganese oxide (3b502)<br>100 mg Zn, zinc oxide (3b603)<br>0.21 mg I, calcium iodate anhydrate (3b202)<br>0.30 mg Se, sodium selenite (3b801)<br>Enzymes:<br>500 FYT 6-phytase (3.1.3.26) (4a18) |

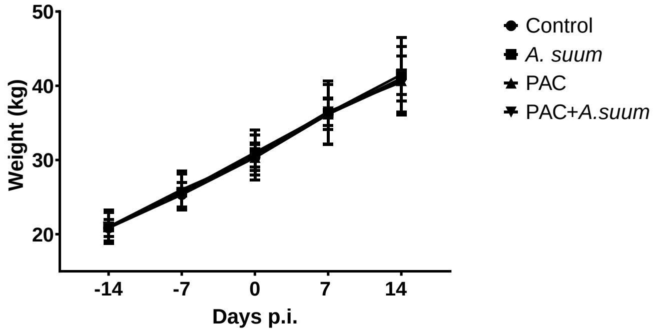


**Figure S2 Pig bodyweights**

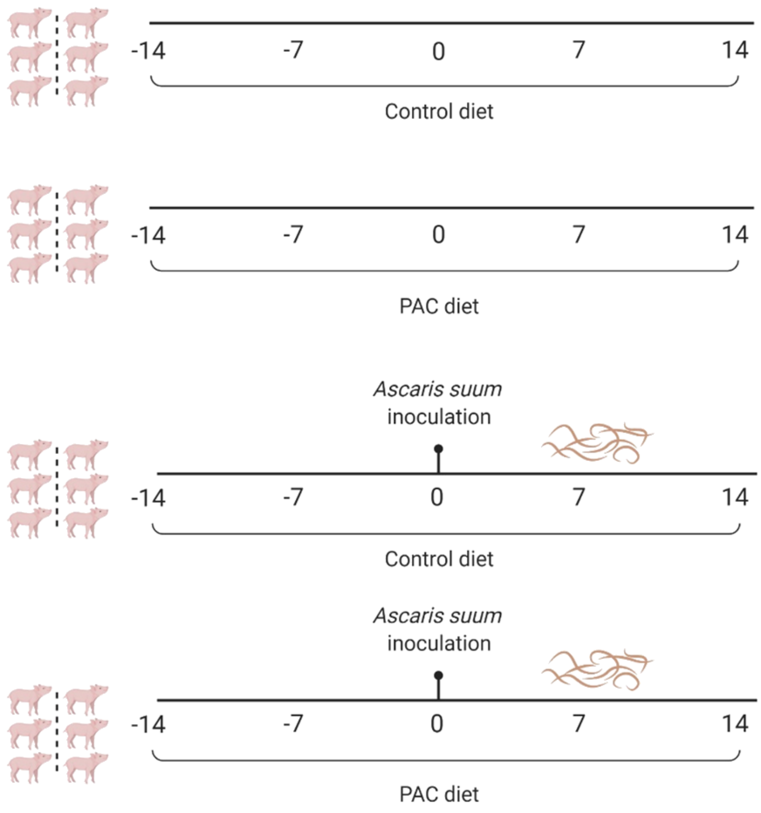


**Figure S3 Experimental study design**

Figure created with Biorender.com

**Table S2 UMI containing primers used for 16S rRNA gene amplification.**

| Primers | Primer Sequence |
|---|---|
| UMI_338Fa | 5'- GTCTCGTGGGCTCGG- NNNNNNNNNNNNNNN - ACWCCTACGGGWGGCAGCAG-3' |
| UMI_338Fb | 5'- GTCTCGTGGGCTCGG- NNNNNNNNNNNNNNN - GACTCCTACGGGAGGCWGCAG-3' |
| UMI_27Fa | 5'- GTCTCGTGGGCTCGG- NNNNNNNNNNNNNNN - AGAGTTTGATYMTGGCTYAG-3' |
| UMI_27Fb | 5'- GTCTCGTGGGCTCGG- NNNNNNNNNNNNNNN - AGGGTTCGATTCTGGCTCAG-3' |
| UMI_1540R | 5'- GTCTCGTGGGCTCGG- NNNNNNNNNNNNNNN - TACGGYTACCTTGTTACGACT-3' |
| UMI_1391R | 5'- GTCTCGTGGGCTCGG- NNNNNNNNNNNNNNN - GACGGGCGGTGTGTRCA-3' |

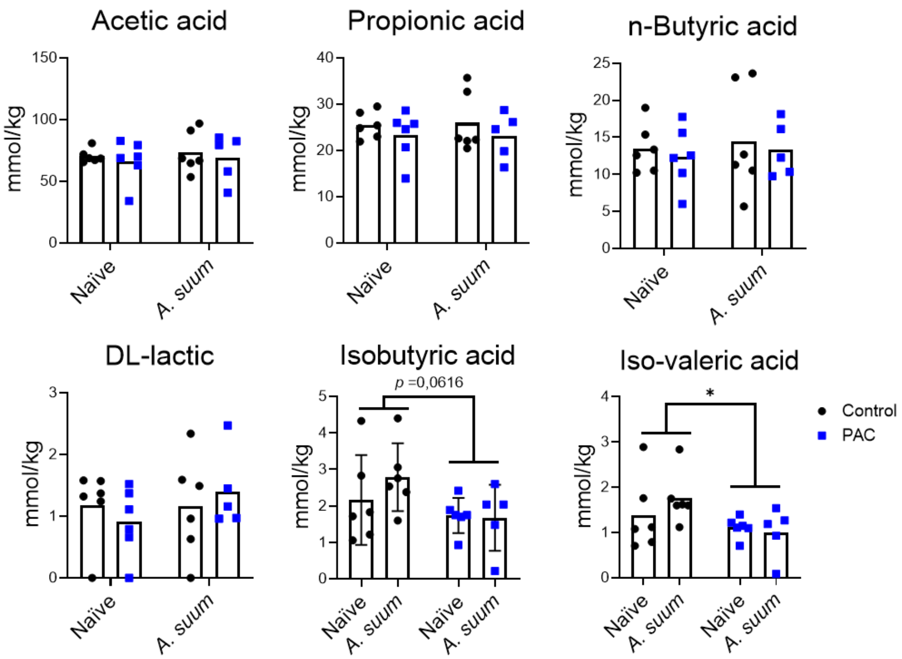


**Figure S3 Short chain and branched chain fatty acid concentrations in colon samples**
